# Haplotype-resolved and integrated genome analysis of the cancer cell line HepG2

**DOI:** 10.1101/378497

**Authors:** Bo Zhou, Steve S. Ho, Stephanie U. Greer, Noah Spies, John M. Bell, Xianglong Zhang, Xiaowei Zhu, Joseph G. Arthur, Seunggyu Byeon, Reenal Pattni, Ishan Saha, Yiling Huang, Giltae Song, Dimitri Perrin, Wing H. Wong, Hanlee P. Ji, Alexej Abyzov, Alexander E. Urban

## Abstract

The HepG2 cancer cell line is one of the most widely-used biomedical research and one of the main cell lines of ENCODE. Vast numbers of functional genomics and epigenomics datasets have been produced to characterize its biology. However, the correct interpretation such data requires an understanding of the cell line’s genome sequence and genome structure. Using a variety of sequencing and analysis methods, we identified a wide spectrum of HepG2 genome characteristics: copy numbers of chromosomal segments, SNVs and Indels (corrected for aneuploidy), phased haplotypes extending to entire chromosome arms, loss of heterozygosity, retrotransposon insertions, structural variants (SVs) including complex and somatic genomic rearrangements. We also identified allele-specific expression and DNA methylation genome-wide and assembled an allele-specific CRISPR/Cas9 targeting map.

**SIGNIFICANCE:** Haplotype-resolved and comprehensive whole-genome analysis of a widely-used cell line for cancer research and ENCODE, HepG2, serves as an essential resource for unlocking complex cancer gene regulation using a genome-integrated framework and also provides genomic context for the analysis of ~1,000 functional datasets to date on ENCODE for biological discovery. We also demonstrate how deeper insights into genomic regulatory complexity are gained by adopting a genome-integrated framework.

## INTRODUCTION

Genomic instability is a hallmark of cancer where critical genomic changes create gene fusions, the disruption of tumor-suppressor, and the amplification of oncogenes (Adey et al., 2013; Hanahan and Weinberg, 2011; Negrini et al., 2010). A comprehensive knowledge of the mutations and larger structural changes that underlie a cancer genome is not only critical for a deeper understanding of the biological processes that drive tumor progression and evolution but also for the development of targeted cancer therapies. The HepG2 cell line is one of the most widely used cancer cell lines used in many areas biomedical research due to its extreme versatility, contributing to over 23,000 publications to date, even more than K562. It is a hepatoblastoma cell line derived from a 15-year-old Caucasian male (Aden et al., 1979; López-Terrada et al., 2009a). Representing the human endodermal lineage, HepG2 cells are widely used as models for human toxicology studies (Dias da Silva et al., 2013; Kamalian et al., 2015; Menezes et al., 2013; Sahu et al., 2012; Schoonen et al., 2005), including toxicogenomic screens using CRISPR-Cas9 (Xia et al., 2016), in addition to studies on drug metabolism (Alzeer and Ellis, 2014), cancer (Xu et al., 2013), liver disease (Hao et al., 2014), gene regulatory mechanisms (Huan et al., 2014), and biomarker discovery (Mangrum et al., 2015). As one of the main cell lines of the ENCyclopedia Of DNA Elements Project (ENCODE), HepG2 has been used to generate close to 1,000 datasets for ENCODE (Sloan et al., 2016).

The functional genomic and epigenomics aspects of HepG2 cells have been extensively studied with approximately 325 ChIP-Seq, 300 RNA-Seq, and 180 eCLIP datasets available through ENCODE in addition to recent single-cell methylome and transcriptome datasets (Hou et al., 2016). However, the genome sequence and higher-order genomic structural features of HepG2 have never been characterized in a comprehensive manner, even though the HepG2 cell line has been known to contain multiple chromosomal abnormalities (Chen et al., 1993; Simon et al., 1982). As a result, the extensive HepG2 functional genomics and epigenomics studies conducted to date were done without reliable genomic contexts for accurate interpretation.

Here, we report the first global, integrated, and haplotype-resolved whole-genome characterization of the HepG2 cancer genome that includes copy numbers (CN) of large chromosomal regions at high-resolution, single-nucleotide variants (SNVs, also including single-nucleotide polymorphisms, i.e. SNPs) and small insertions and deletions (indels) with allele-frequencies corrected by CN in aneuploid regions, loss of heterozygosity, mega-base-scale phased haplotypes, and structural variants (SVs), many of which are haplotype-resolved (Figure 1, Figure S1). The datasets generated in this study form an integrated, phased, high-fidelity genomic resource that can provide the proper contexts for future experiments that rely on HepG2’s unique characteristics. We show how knowledge about HepG2’s genomic sequence and structural variations can enhance the interpretation of functional genomics and epigenomics data. For example, we integrated HepG2 RNA-Seq data and whole-genome bisulfite sequencing data with ploidy and phasing information and identified many cases of allele-specific gene expression and allele-specific DNA methylation. We also compiled a phased CRISPR map of loci suitable for allele specific-targeted genome editing or screening. Finally, we demonstrate the power of this resource by providing compelling insights into the mutational history of HepG2 and oncogene regulatory complexity derived from our datasets. The technical framework demonstrated in this study is also suitable for the study of other cancer cell lines and primary tumor samples.

**Figure 1.**
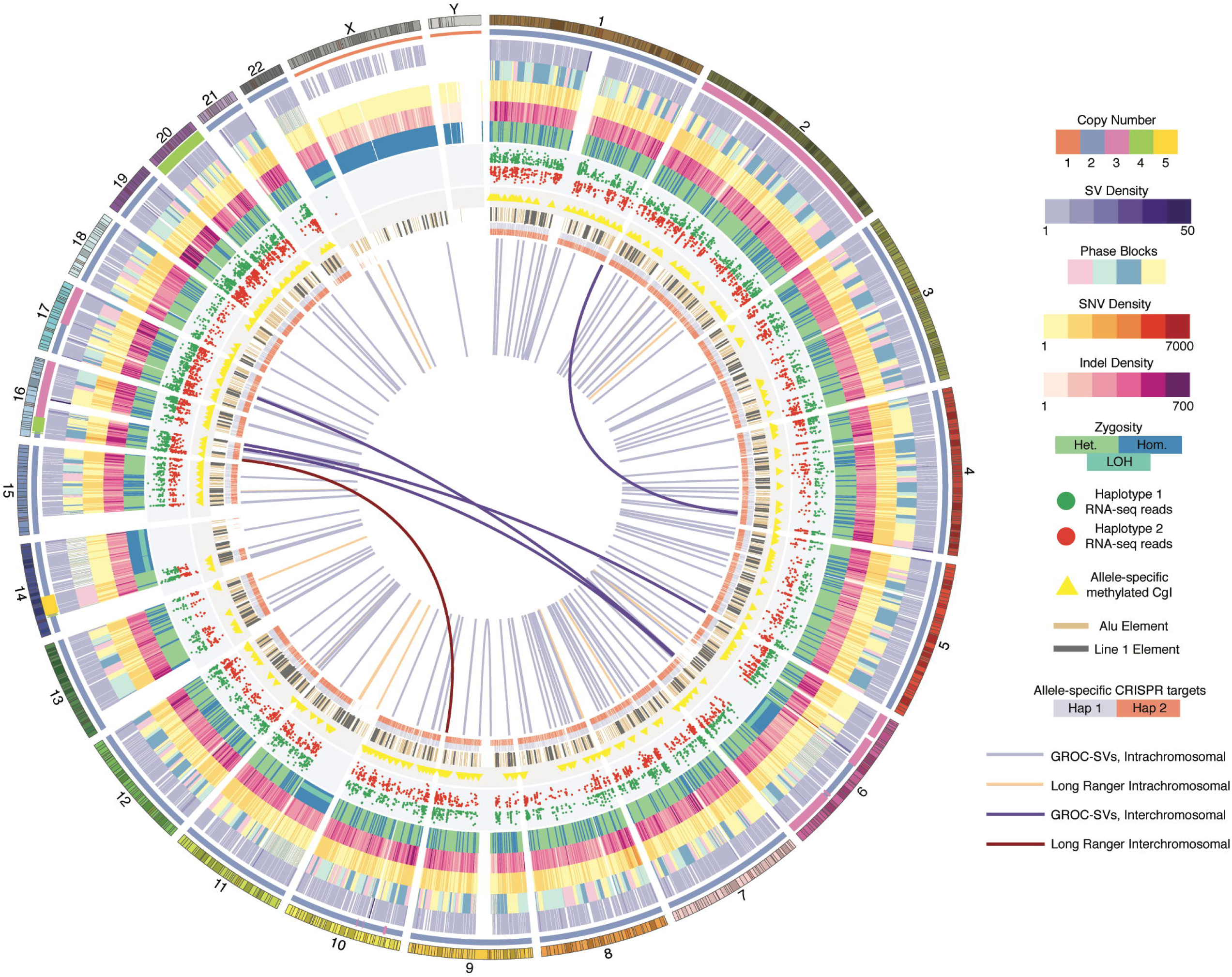
Comprehensive Overview of the HepG2 Genome. Circos visualization (Krzywinski et al. 2009) of HepG2 genome variants with the following tracks in concentric order starting with outermost “ring”: human genome reference track (hg19); large CN changes (colors correspond to different CN, see legend panel); in 1.5 Mb windows, merged SV density (deletions, duplications, inversions) called using BreakDancer, BreakSeq, PINDEL, LUMPY, and Long Ranger; phased haplotype blocks (demarcated with 4 colors for clearer visualization); SNV density in 1 Mb windows; Indel density in 1 Mb windows; dominant zygosity (heterozygous or homozygous > 50%) in 1 Mb windows; regions with loss of heterozygosity; allele-specific expression; CpG islands exhibiting allele-specific DNA methylation; non-reference LINE1 and Alu insertions; allele-specific CRISPR target sites; large-scale SVs resolved by using Long Ranger (peach: intrachromosomal: dark maroon: interchromosomal); by using GROC-SVs (light-purple: intrachromsomal; dark-purple: interchromosomal).

## RESULTS

### Karyotyping

We obtained HepG2 cells from the Stanford ENCODE Production Center. The cells exhibit a hyperdiploid karyotype of 49 to 52 chromosomes (Figure 2A). Of the 20 metaphase HepG2 cells analyzed using the GTW banding method, 15 of cells were complexly and variably abnormal and also characterized by multiple structural and numerical abnormalities. These include translocation between the chromosome 1p and 21p, trisomies of chromosomes 2, 16, and 17, tetrasomy of chromosome 20, uncharacterized arrangements of chromosomes 16 and 17, and a variable number of marker chromosomes. Five cells demonstrated greater than 100 chromosomes and represent a tetraploid expansion of the stemline described. This tetraploid expansion is consistent with previously published results (Simon et al. 1982) but also absent from other published cytogenetic analyses of HepG2 (Chen et al. 1993), suggesting the clonal evolution arose during tumor-genesis or early in the establishment of the HepG2 cell line. Although the ploidies of all chromosomes in our HepG2 cell line were supported by previous published karyotypes (Chen et al., 1993; Simon et al., 1982; Wong et al., 2000), variations do exist and also among the various published analyses especially for chromosomes 16 and 17, suggesting that karyotypic differences exist between different HepG2 cell lines (Table S1).

**Figure 2.**
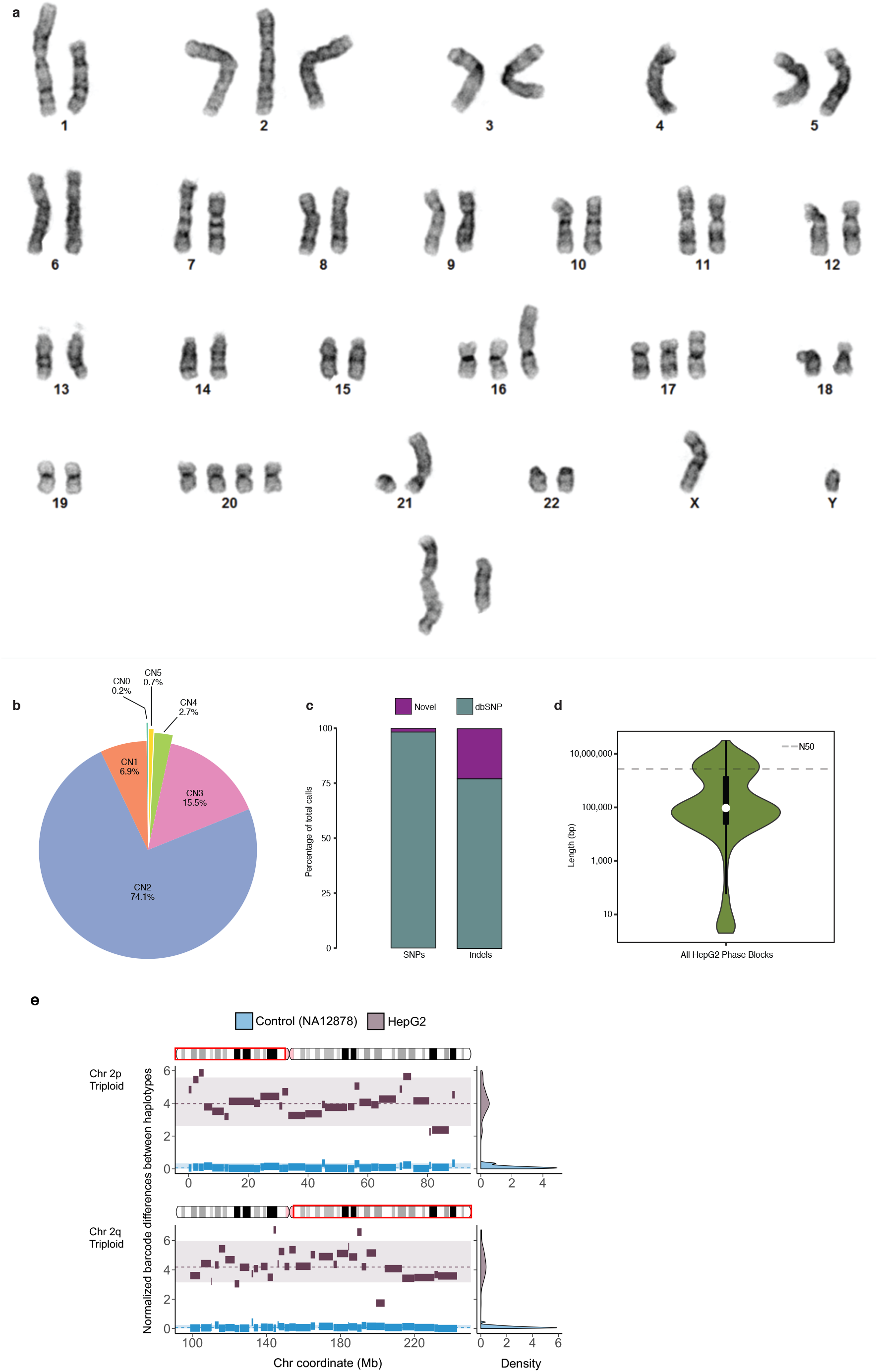
HepG2 Karyogram and Callset Overview. (A) Representative karyogram of HepG2 cells by GTW banding which shows multiple numerical and structural abnormalities including a translocation between the short arms of chromosomes 1 and 21, trisomies of chromosomes 12, 16, and 17, tetrasomy of chromosome 20, uncharacterized rearrangements of chromosomes 16 and 17 and a two marker chromosomes. ISCN 2013 description: 49~52,XY,t(1:21)(p22;p11),+2,+16,add(16)(p13),?+17,?add(17)(p11.2),+20,+20,+1~3mar[cp15]/ 101~106,idemx2[cp5]. (B) CNs (by percentage) across the HepG2 genome. (C) Percentage of HepG2 SNVs and Indels that are novel and known (in dbSNP). (D) Violin plot with overlaid boxplot of phased haplotype block sizes, with N50 represented as a dashed line (N50 = 6,792,324 bp) with log-scaled Y-axis. (E) X-axis: chromosome coordinate (Mb). Y-axis: difference in unique linked-read barcode counts between major and minor haplotypes, normalized by SNV density. Haplotype blocks from of normal control sample (NA12878) in blue and from HepG2 in dark gray. Density plots on the right reflects the distribution of the differences in haplotype-specific barcode counts for control sample HepG2. Significant difference (one-sided t-test, p<0.001) in haplotype-specific barcode counts indicate aneuploidy and haplotype imbalance. Haplotype blocks (with ≥100 phased SNVs) generated from Long Ranger (Dataset 2) for the major and minor haplotypes were then “stitched” to mega-haplotypes encompassing the entire triploid chromosome arms of 2p and 2q.

### High-Resolution Ploidy Changes in HepG2

To obtain a high-resolution aneuploid map i.e. large CN changes by chromosomal region in HepG2, WGS coverage across the genome was first calculated in 10 kb bins and plotted against percent GC content where four distinct clusters were clearly observed (Abyzov et al., 2011) (Figure S2). CNs were assigned to each cluster based on the ratio between its mean coverage and that of the lowest cluster (CN=1). These assigned large CN changes by chromosomal region confirm the hyperdiploid state of the HepG2 genome as identified by karyotyping (Figure 2A, Figure S2, Table S2). We see that 74.1% of the HepG2 genome has a baseline copy number of two (consistent with karyotype), 15.5% copy number of three, 2.7% copy number of four, 0.7% has a copy number of five, and 6.9% in a haploid state (Figure 2B, Table S2). Furthermore, these high-resolution CN changes across the HepG2 genome were also confirmed by two independent replicates of Illumina Infinium Multi-Ethnic Global-8 arrays (MEGA) array data (Figure S3A, Supplementary Data). We found increased CN (CN=3) over the oncogene *VEGFA* (6p21.1), which was found to be recurrently duplicated in cases of hepatocellular carcinoma (Cancer Genome Atlas Research Network, 2017).

### SNVs and indels

We identified SNVs and indels in HepG2 by taking into account the CN of the chromosomal regions in which they reside so that heterozygous allele frequencies can be assigned accordingly (e.g. 0.33 and 0.67 in triploid regions; 0.25, 0.50, and 0.75 in tetraploid regions). Using GATK Haplotypecaller (McKenna et al., 2010), we identified a total of ~3.34M SNVs (1.90M heterozygous, 1.44M homozygous) and 0.90M indels (0.60M heterozygous, 0.29M homozygous) (Table 1, Dataset 1). Interestingly, there are 12,375 heterozygous SNVs and indels that have more than two haplotypes in chromosomal regions with CN>2 (Dataset 1). In addition, chromosome 22 and large continuous stretches of chromosomes 6, 11, and 14 show striking loss of heterozygosity (LOH) (Figure 1, Table S3). Since genomic data from healthy tissue that correspond to HepG2 cells is not available, we intersected these SNVs and indels with dbsnp138 (Sherry et al., 2001) and found the overlapping proportion to be ~98% and ~78% respectively (Figure 2C). This suggests that HepG2 has accumulated a significant number of SNVs and indels relative to inherited. We found that 377 SNVs and 255 indels are private protein-altering (PPA) after filtering out those that overlapped with The 1000 Genomes Project (The 1000 Genomes Project Consortium et al., 2015) or the Exome Sequencing Project (Fu et al., 2012) (Table 1, Table S4). Moreover, the intersection between the filtered PPA variants and the Catalogue of Somatic Mutations in Cancer (COSMIC) is 39% and 16% for SNVs and indels respectively (Table S5). The gene overlap between HepG2 PPA and the Sanger Cancer Gene Census is 19 (Table S6). HepG2 PPA variants include oncogenes and tumor suppressors such as *NRAS* (Pylayeva-Gupta et al., 2011)*, STK11/LKB1* (Zhou et al., 2014), and *PREX2* (Berger et al., 2012; Yang et al., 2016) as well as other genes recently found to play critical roles in driving cancer such as *CDK12* (Paculová and Kohoutek, 2017) and *IKBKB* (Kai et al., 2014; Xia et al., 2012)*. RP1L1*, which was recently found to be significantly mutated in hepatocellular carcinoma (Cancer Genome Atlas Research Network 2017) is also present among the PPA variants.

**Table 1.**
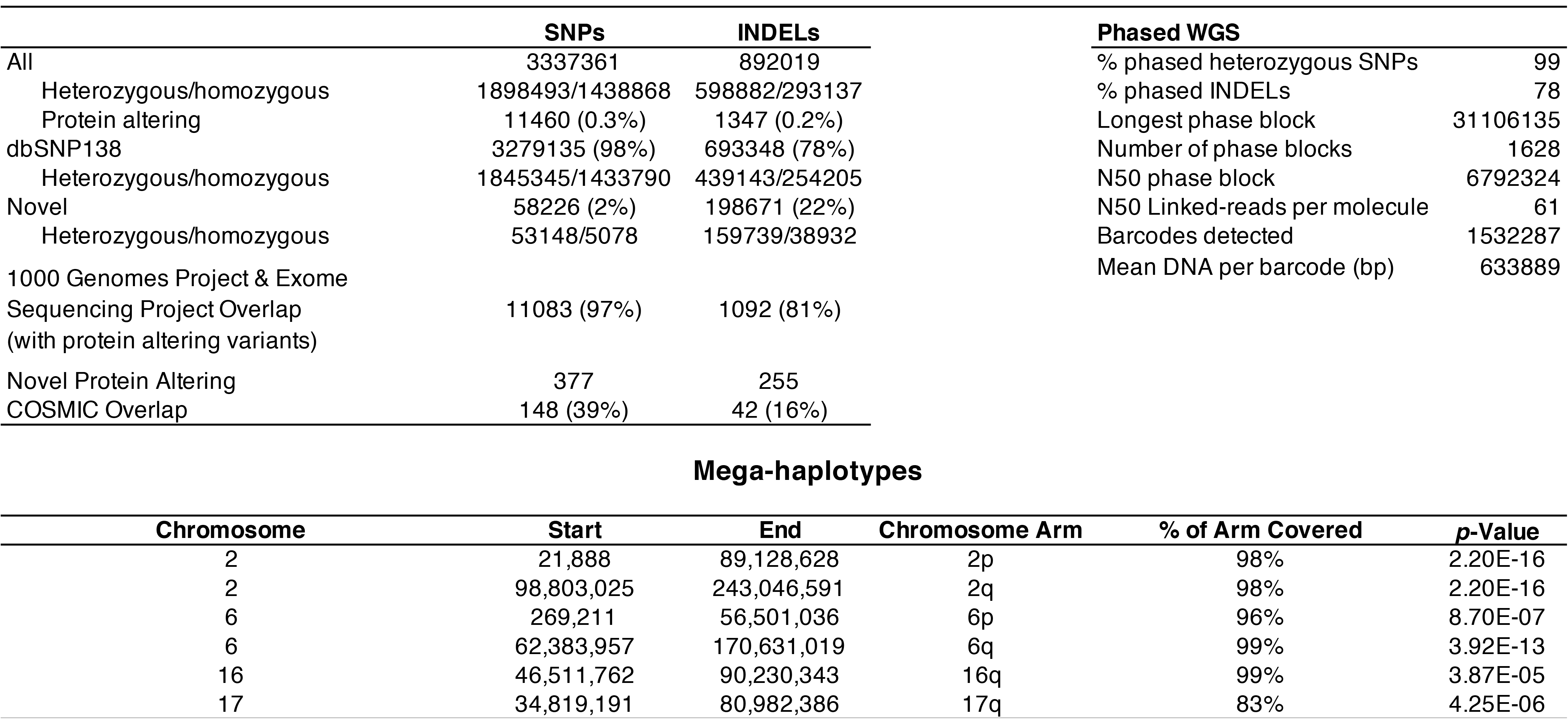
Summary of HepG2 Small Variant Calls and Phasing Results.

### Resolving Haplotypes

We phased the heterozygous SNVs and indels in the HepG2 genome by performing 10X Genomics Chromium linked-read library preparation and sequencing (Marks et al., 2018; Zheng et al., 2016). Post sequencing quality control analysis show that 1.49 ng or approximately 447 genomic equivalents of high molecular weight (HMW) genomic DNA fragments (mean=68 kb, 96.1% > 20 kb, 22.0% >100 kb) were partitioned into 1.53 million oil droplets and uniquely barcoded (16 bp). This library was sequenced (2×151 bp) to 67x genome coverage with half of all reads coming from HMW DNA molecules with at least 61 linked reads (N50 Linked-Reads per Molecule) (Table 1). We estimate the actual physical coverage (C_F_) to be 247x. Coverage of the mean insert by sequencing (C_R_) is 18,176 bp (284 bp X 64) or 30.8%, thus the overall sequencing coverage C = C_R_ X C_F_ = 67x. Distributed over 1628 haplotype blocks (Table 1, Dataset 2), 1.87M (98.7%) of heterozygous SNVs and 0.67M (77.9%) of indels in HepG2 were successfully phased. The longest phased haplotype block is 31.1Mbp (N50=6.80Mbp) (Figure 2D, Table 1, Dataset 2); however, haplotype block lengths vary widely across different chromosomes (Figure S4, Figure 1). Poorly phased regions correspond to regions exhibiting LOH (Table S3, Figure 1, Dataset 2).

### Construction of Mega-Haplotypes of Entire Chromosome Arms

We constructed mega-haplotypes of entire chromosome arms by leveraging the haplotype imbalance in aneuploid regions in the HepG2 genome where phased haplotype blocks derived from linked-reads were “stitched” together (Table 1, Figure 2E, Supplementary Data). Briefly, by using a recently developed method (Bell et al., 2017) specifically for cancer genomes, we counted linked-read barcodes for each phased heterozygous SNVs in haplotype blocks with ≥100 phased SNVs (Dataset 2). Because each barcode is specific for a HMW DNA molecule, the total number of unique barcodes is directly correlated with the number of individual HMW DNA molecules that were sequenced. The fractional representation of a particular genomic sequence (or locus) can be obtained by counting the total number of unique barcodes associated with that particular genomic sequence. Consequently, for each phased haplotype with CN>2, major and minor haplotypes can be assigned according to the number of barcodes associated with each haplotype (Figure 2E), where the major haplotype is the haplotype with more associated unique barcodes. In genomic regions where CN=2, the two haplotypes are expected to have similar numbers of unique barcodes. In this method (Bell et al., 2017), a matched control for comparison is required to confidently discriminate between the major and minor haplotypes. Here, we used NA12878 as normal control because no matching normal tissue sample is available for HepG2 (Figure 2E). After performing the normalization procedures and statistical tests described in (Bell et al., 2017) to verify haplotype imbalance or aneuploidy genomic regions in HepG2, we then “stitched” together contiguous blocks of phased major and minor haplotypes respectively. Using this approach, a total of 6 autosomal mega-haplotypes were constructed (Table 1, Supplementary Data); 4 of which encompass entire (or >96%) chromosome arms: 2p, 2q, 6p, and 16q (Fig. 3). The largest mega-haplotype is approximately 144 Mb long (2q).

**Figure 3.**
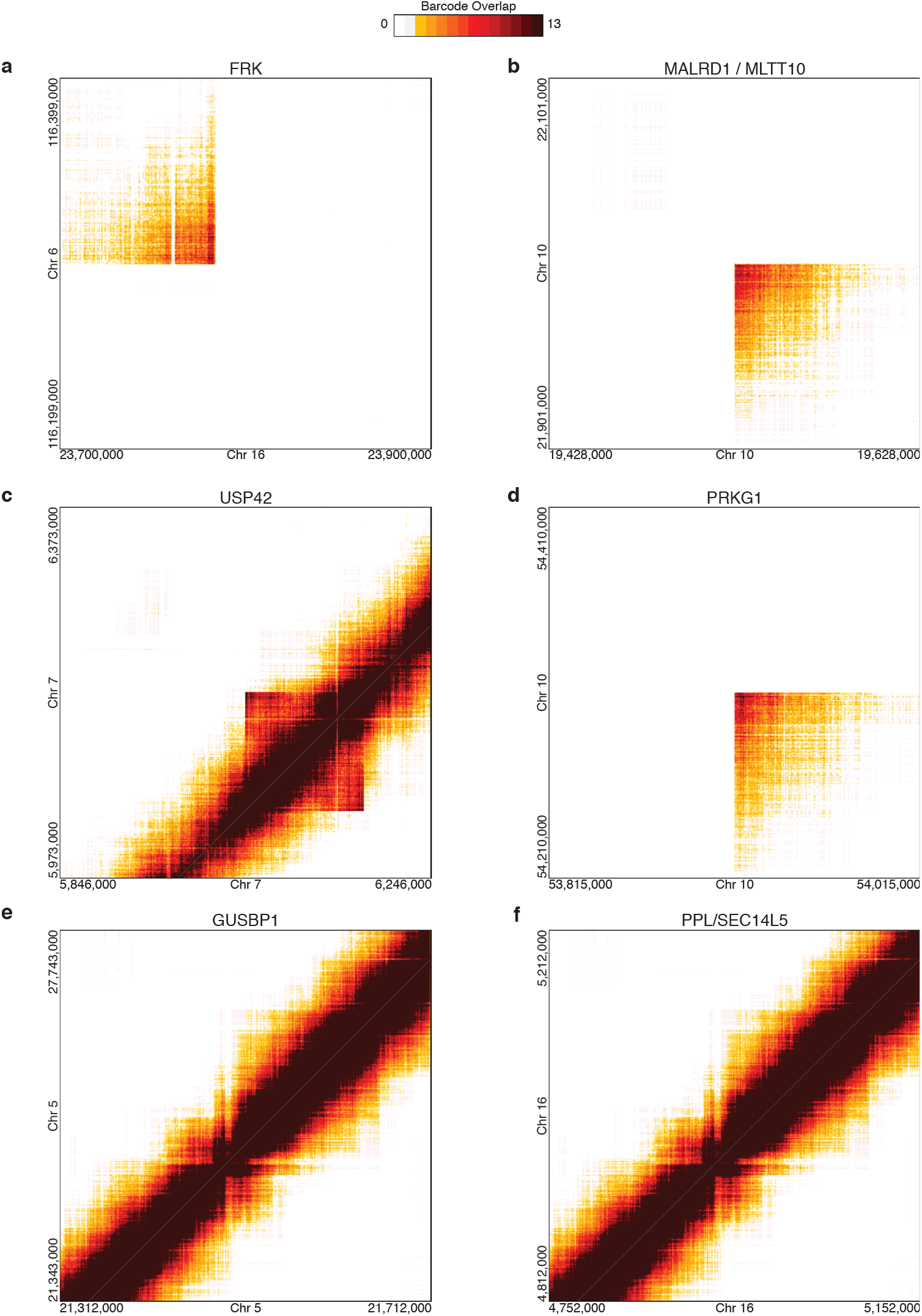
Large SVs in HepG2 Resolved from Linked-Read Sequencing using Long Ranger. HepG2 SVs resolved by identifying identical linked-read barcodes in distant genomic regions with non-expected barcode overlap for identified using Long Ranger (Marks et al., 2018; Zheng et al., 2016). (A) Disruption of *FRK* by translocation between chromosomes 6 and 16. (B) 2.47 Mb intra-chromosomal rearrangement between *MALRD1* and *MLLT10* on chromosome 10. (C) 127 kb duplication on chromosome 7 resulting in partial duplications of *USP42* and *PMS2.* (D) 395 kb duplication within *PRKG1* on chromosome 10. (D) 194 kb deletion within *PDE4D* on chromosome 5 (E) 286 kb deletion within *AUTS2* on chromosome 7 (E) 31.3 kb inversion within *GUSBP1* on chromosome 5. (F) 60.4 kb inversion that disrupts *PPL* and *SEC14L5*.

### Using Linked-Reads to Identify and Reconstruct Large and Complex SVs

From the linked reads, breakpoints of large-scale SVs can be identified by searching for distant genomic regions with linked-reads that have large numbers of overlapping barcodes. SVs can also be assigned to specific haplotypes if the breakpoint-supporting reads contain phased SNVs or indels (Marks et al., 2018; Zheng et al., 2016). Using this approach (implemented by the Long Ranger software from 10X Genomics), we identified 97 large SVs >30 kb (99% phased) (Dataset 3) and 3,473 deletions between 50 bp and 30 kb (78% phased) (Dataset 4). The large SVs include inter- and intra-chromosomal rearrangements (Figure 3A, B), duplications (Figure 3C, D), deletions (Figure 3E, F), and inversions (Figure 3G, H). A remarkable example is the haplotype-resolved translocation between chromosomes 16 and 6 (Figure 3A) resulting in the disruption of the non-receptor Fyn-related tyrosine kinase gene *FRK*, which has been identified as a tumor suppressor (Brauer and Tyner, 2009; Yim et al., 2009). Another example is the 127 kb tandem duplication on chromosome 7 (Figure 3C) that results in the partial duplication of genes *PMS2*, encoding a mismatch repair endonuclease, and *USP42*, encoding the ubiquitin-specific protease 42. An interesting large SV is the 395 kb duplication within *PRKG1* (Figure 3D), which encodes the soluble l-alpha and l-beta isoforms of cyclic GMP-dependent protein kinase. We also identified a 193 kb homozygous deletion in *PDE4D* for HepG2 using linked-read sequencing where six internal exons within the gene are deleted (Figure 3D).

Furthermore, we also used the long-range information from the deep linked-reads sequencing dataset to identify, assemble, and reconstruct the breakpoints of SVs in the HepG2 genome using a recently developed method called Genome-wide Reconstruction of Complex Structural Variants (GROC-SVs) (Spies et al., 2017). Here, HMW DNA fragments that span breakpoints are statistically inferred and refined by quantifying the barcode similarity of linked-reads between pairs of genomic regions similar to Long Ranger (Zheng et al., 2016). Sequence reconstruction is then achieved by assembling the relevant linked reads around the identified breakpoints from which SVs are automatically reconstructed. Breakpoints that also have supporting evidence from the 3kb-mate pair dataset (see Methods) are indicated as high-confidence events. GROC-SVs called a total of 140 high-confidence breakpoints including 4 inter-chromosomal events (Figure 1, Dataset 5, Figure 4A-D); 138 of the breakpoints were successfully sequence-assembled with nucleotide-level resolution of breakpoints as well the exact sequence in the cases where nucleotides have been added or deleted. We identified striking examples of inter-chromosomal rearrangements or translocations in HepG2 between chromosomes 1 and 4 (Figure 4A) and between chromosomes 6 and 17 (Figure 4B) as well as breakpoint-assembled large genomic deletions (Figure 4C, Dataset 5). This break-point assembled 335 kb heterozygous deletion is within the *NEDD4L* on chromosome 18. Lastly, we identified a large (1.3 mb) intra-chromosomal rearrangement that deletes large portions of *RBFOX1* and *RP11420N32* in one haplotype on chromosome 16 using deep linked-read sequencing (Figure 4D, Dataset 3, Dataset 5).

**Figure 4.**
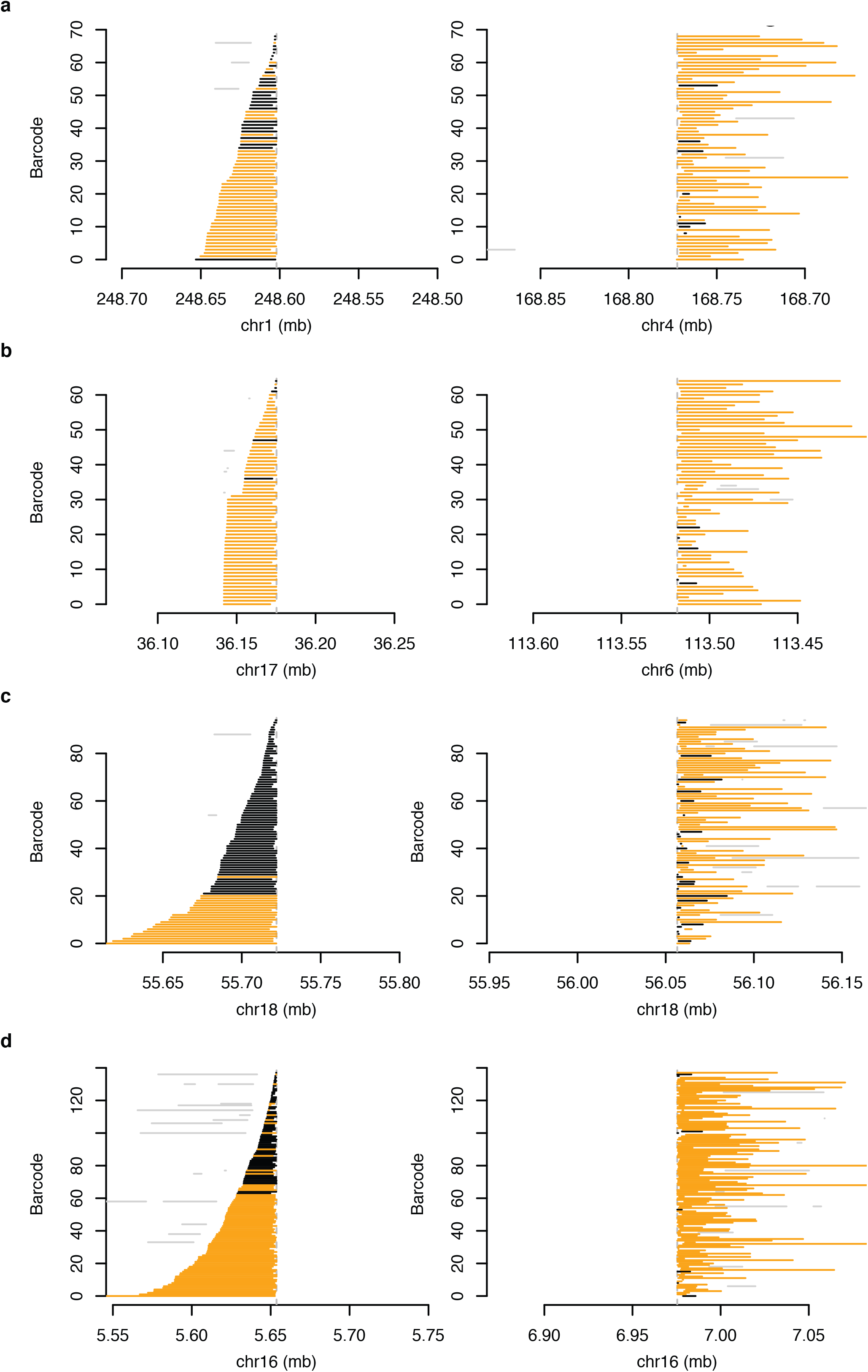
HepG2 SVs Reconstructed and Assembled Using GROC-SVs in HepG2. (A-D) Each line depicts a fragment inferred from 10X-Genomics data based on clustering of reads with identical barcodes (Y-axis) identified from GROC-SVs (Spies et al., 2017). Abrupt ending (dashed vertical line) of fragments indicates location of SV breakpoint. All breakpoints depicted are validated by 3 kb-mate-pair sequencing data. Fragments are phased locally with respect to surrounding SNVs (haplotype-specific) are in orange, and black when no informative SNVs are found nearby. Gray lines indicate portions of fragments that do not support the current breakpoint. (A) Translocation between chromosomes 1 and 4. Linked-read fragments containing overlapping barcodes that map to chromosome 1 end abruptly near 248.60 mb indicating a breakpoint, and then continues simultaneously near 168.75 mb on chromosome 4. (B) Translocation between chromosomes 6 and 17. Linked-read fragments containing overlapping barcodes that map to chromosome 17 end abruptly near 36.17 mb indicating a breakpoint and then continues simultaneously near 113.52 mb on chromosome 6. (C) Large (335 kb) heterozygous deletion within *NEDD4L* on chromosome 18. (D) Large (1.3 mb) intra-chromosomal rearrangement that deletes large portions of *RBFOX1* and *RP11420N32* on chromosome 16.

We then employed “gemtools” (Greer et al., 2017) to resolve and phase large and complex SVs in the HepG2 genome. We identified a complex SV on chromosome 8 that involves a small deletion downstream of *ADAM2* which is also within a larger tandem duplication leading to the amplification of the oncogene *IDO1* (Platten et al., 2015) and the first half of *IDO2* (Figure 5). Two allele-specific deletions 700 kb and 200 kb respectively were identified in the *PDE4D* on chromosome 5 (Figure 5). Since chromosome 5 is triploid in HepG2 (Figure 1, Figure 2A, Supplementary Information), we see approximately twice as much linked-reads barcode representation for the allele harboring the 200 kb deletion, suggesting that this allele of *PDE4D* has two copies and the allele harboring the 700 kb deletion has one copy (Figure 5). Similarly, we also identified two allele-specific deletions, 290 kb and 160 kb respectively within *AUTS2* on chromosome 7 (Figure 5). Interestingly, for the allele harboring the 160 kb deletion, the non-deleted reference allele is also present at much larger frequency as indicted by the total number of linked-read barcodes, suggesting that the allele harboring the 160 kb deletion within *AUTS2* occurs in a fraction of HepG2 cells or sub-clonally (Figure 5). Estimating from the total number of linked-read barcodes that are associated with this 160 kb allele-specific deletion in *AUTS2*, we estimate that this deletion occur at a frequency of ~10%. In other words, 10% of HepG2 cells have both deletions rendering a large portion of *AUTS2* in HepG2 cells deleted completely in ~10% of HepG2 cells whereas the other 90% of HepG2 cells carry only the 290 kb deletion within *AUTS2*. All breakpoints identified using “gemtools” were individual PCR and Sanger sequencing verified (Table S7).

**Figure 5.**
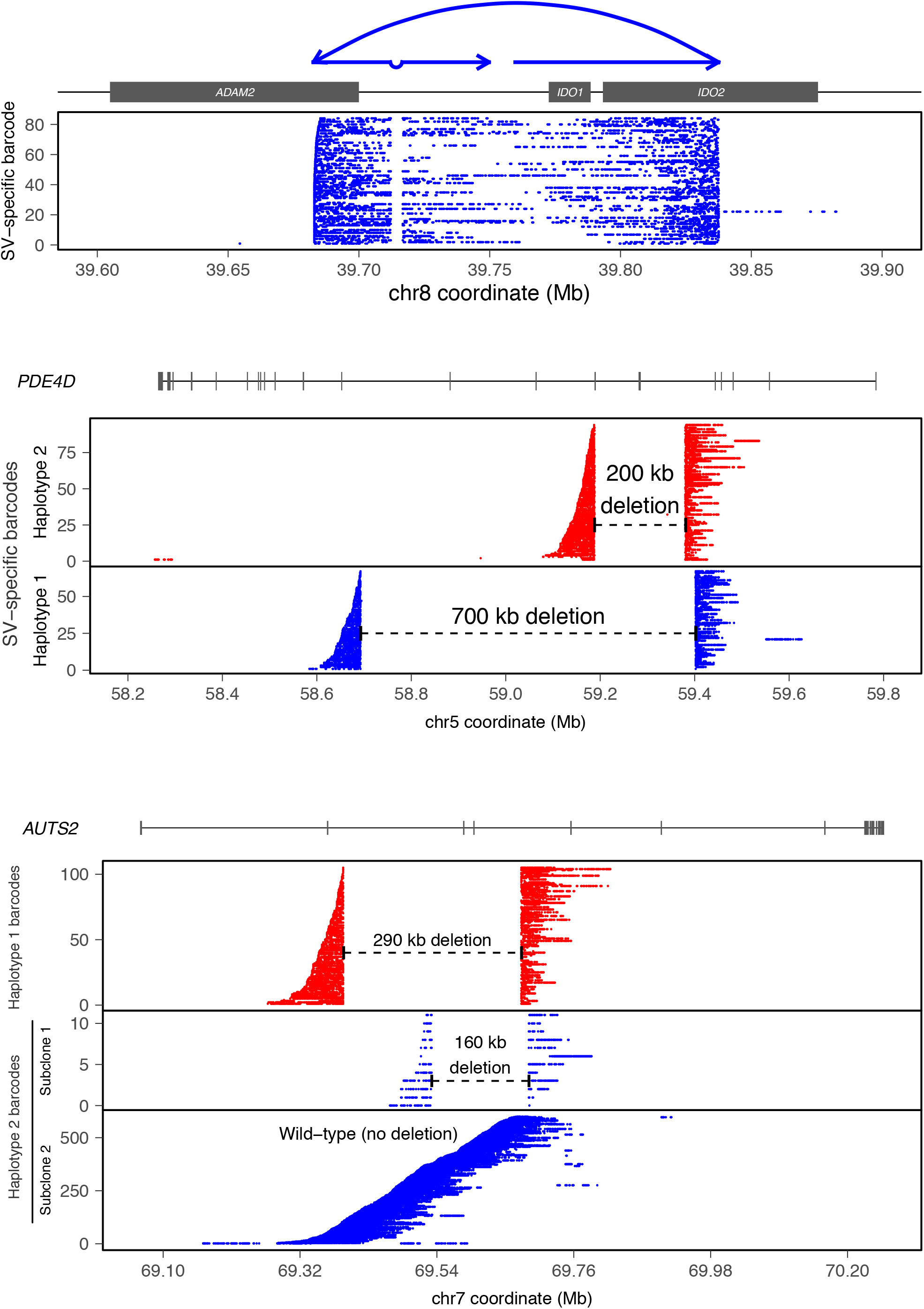
Large and complex haplotype-resolved SVs using gemtools (Greer et al., 2017). Each SV is identified from linked-reads clustered by identical barcodes (i.e. SV-specific barcodes, Y-axis) indicative of single HMW DNA molecules (depicted by each row) that span SV breakpoints. Haplotype-specific SVs are represented in blue and red. X-axis: hg19 genomic coordinate. (Top) Complex SV on chromosome 8 involving a 4585 bp deletion downstream of *ADAM2*. This deletion is within a tandem duplication leading to the amplification of the *IDO1* and the first half of *IDO2*. The presence of HMW molecules sharing the same linked-read barcodes spanning both breakpoints indicates a *cis* orientation and occurrence on only one allele of this locus. Schematic diagram of the rearranged structures drawn above the plot. (Middle) Two haplotype-resolved deletions 700 kb (blue) and 200 kb (red) respectively occurring on two separate alleles within of *PDE4D* on chromosome 5 – the spanning HMW molecules for each deletion do not share SV-specific barcodes, indicating that these deletions are in *trans*. (C) Two haplotype-resolved deletions, 290 kb (red) and 160 kb (blue) respectively, within *AUTS2* on chromosome 7. The reference allele of *AUTS2* without the deletion (Haplotype 2) is also detected and resolved by linked-reads (blue, bottom panel). The 160 kb deletion on Haplotype 2 occurs sub-clonally.

**Figure 6.**
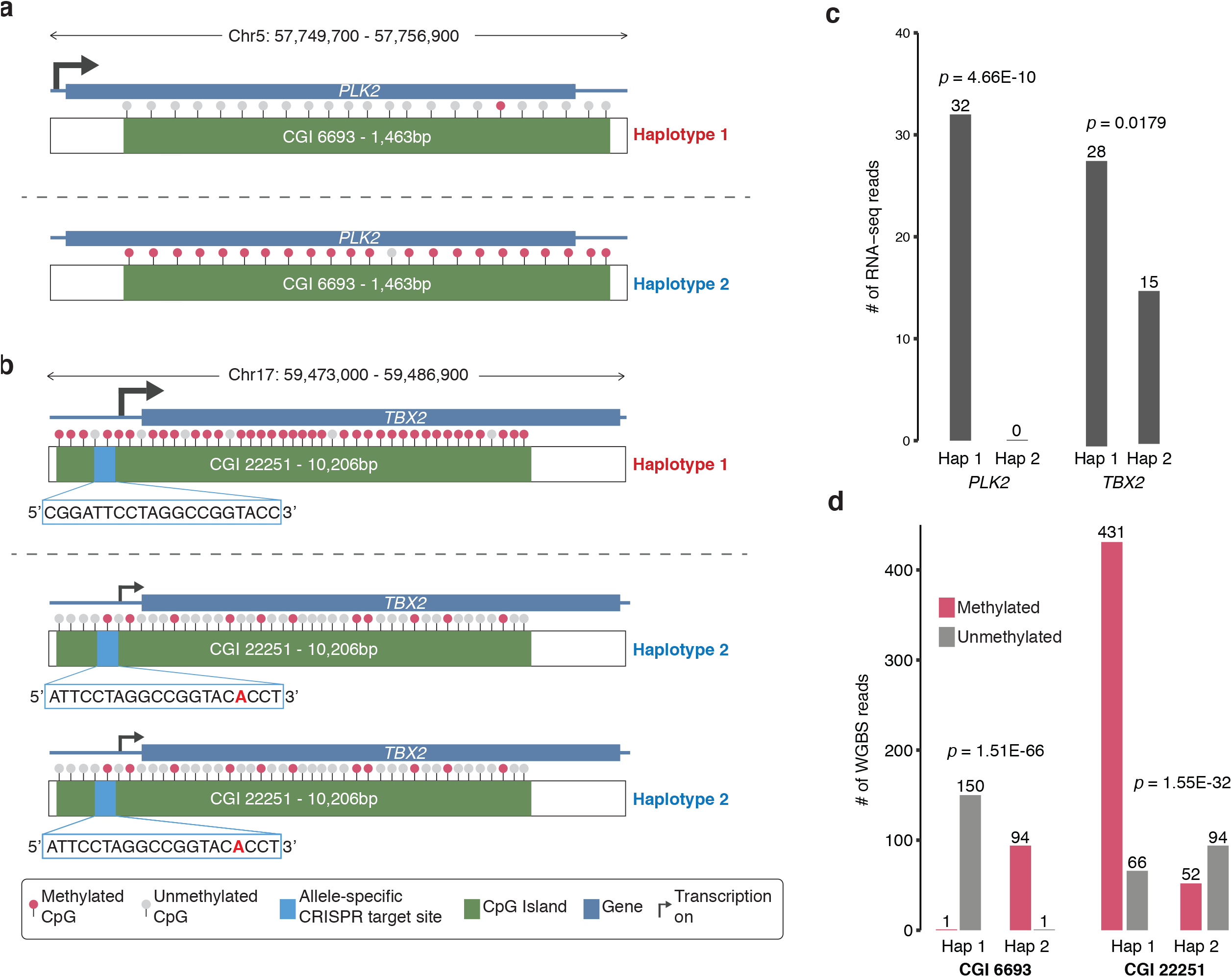
Genomic Sequence and Structural Context Provides Insight into Regulatory Complexity in HepG2. (A) Chr5:57,755,334-57,756,803 locus containing the serine/threonine-protein kinase gene *PLK2* and CGI 6693 (1,463 bp) where phased Haplotype 1 and Haplotype 2. Allele-specific transcription of *PLK2* from Haplotype 2 only. CpGs in CGI 6693 are mostly unmethylated in Haplotype 2 (expressed) and highly methylated in Haplotype 1 (repressed). (B) Chr17:59,473,060-59,483,266 locus (triploid in HepG2) containing T-box transcription factor gene *TBX2* and CpG Island (CGI) 22251 (10,206 bp) where phased Haplotype 2 has two copies and Haplotype 1 has one copy. Allele-specific transcription of *TBX2* from Haplotype 2 only. CpGs in CGI 22251 are unmethylated in Haplotype 1 (repressed) and methylated in Haplotype 2 (expressed). Allele-specific CRISPR targeting site 1937 bp inside the 5’ region of *TBX2* for both Haplotypes. (C) Number of allele-specific RNA-Seq reads in Haplotypes 1 and 2 for *PLK2* and *TBX2* where both genes exhibit allele-specific RNA expression (*p*=0.4.66E-10 and *p*=0.0179 respectively). (D) Number of methylated and unmethylated phased whole-genome bisulfite-sequencing reads for Haplotypes 1 and 2 in CGI 6693 and CGI 22251 where both CGIs exhibit allele-specific DNA methylation (*p*=1.51E-66 and *p*=1.55E-32 respectively).

### SVs from Mate-Pair Sequencing

To obtain increase sensitivity in the detection medium-sized (1 kb-100 kb) SVs in HepG2, we prepared a 3kb-mate pair library and sequenced (2×151 bp) it to a genome coverage of 7.9x after duplicate removal. The sequencing coverage of each 3 kb insert (C_R_) is 302 bp (or 10% of the insert size) which translates to a physical coverage (C_F_) of 79x. Deletions, inversions, and tandem duplications from the mate-pair library were identified from analysis of discordant read pairs and split reads using LUMPY (Layer et al., 2014). Only SVs that are supported by both discordant read-pair and split-read supports were retained. Using this approach, we identified 122 deletions, 41 inversions, and 133 tandem duplications (Dataset 6). Approximately 76% of these SVs are between 1 kb-10 kb, 86% are between 1kb-100kb (9% between 10 kb-100 kb), and 3% are greater than 100 kb (Dataset 6). Twenty SVs (16 deletions and 4 duplications) were randomly selected for experimental validation using PCR and Sanger sequencing in which 15/16 were successfully validated (93.8%) (Table S7).

### SVs identified from deep short-insert WGS

Deletions, inversions, insertions, and tandem duplications, were identified from the HepG2 WGS dataset using Pindel (Ye et al., 2009), BreakDancer (Chen et al., 2009), and BreakSeq (Lam et al., 2010). Since similar categories of SVs were also identified using mate-pair and linked-read sequencing, these SVs were combined with the SVs identified previously using Long Ranger and LUMPY where variations with support from multiple methods and with greater than 50% reciprocal overlap were merged. In total, 6,405 SVs were obtained from all methods that include 5,226 deletions, 245 duplications, 428 inversions, and 494 insertions (only BreakDancer (Chen et al., 2009) was designed to call insertions) (Supplementary Data). A set of deletion (n=27) and tandem duplication calls (n=4) were randomly selected to confirm by PCR and Sanger sequencing, and 30/32 (94%) events were successfully validated (Table S7). Consistent with previous analysis (Lam et al., 2012), deletions show the highest concordance among the various methods of detection compared to duplication and inversion calls (Figure S5). As expected, we detected a 520 bp deletion in exon 3 of the β-catenin (*CTNNB1*) gene (Dataset 4, Supplementary Data), which was previously documented to exist in HepG2 (López-Terrada et al., 2009b). Interestingly, we found no SVs or PPA mutations in the Wnt-pathway gene *CAPRIN2* (Ding et al., 2008), which had been previously reported for hepatoblastoma (Jia et al., 2014).

### Identification of Non-Reference Alu and LINE1 Insertions

From our deep-coverage short-insert WGS data, we also analyzed the HepG2 genome for non-reference LINE1 and Alu retrotransposon insertions using RetroSeq (Keane et al., 2013) with some modifications. These insertions were identified from paired-end reads that have one of the pair mapping to hg19 uniquely and other mapping to an Alu or LINE1 consensus sequence in full or split fashion (see Methods). Retrotransposon insertion events with greater than five supporting reads were categorized as high confidence and retained (Table S8). We identified 1,899 and 351 non-reference Alu and LINE1 insertions in the HepG2 genome respectively (Figure 1). We randomly chose 8 Alu and 10 LINE1 insertions with split-read support for confirmation using PCR and Sanger sequencing where 87.5% and 100% respectively were successfully validated (Table S8).

### Allele-Specific Gene Expression

Due to the abundance of aneuploidy in the HepG2 genome, CN changes of genomic regions should be taken into account when analyzing for allele-specific gene expression in order to reduce false positives and false negatives. Using the heterozygous SNV allele frequencies in HepG2 (Dataset 1), we re-analyzed two replicates of HepG2 ENCODE RNA-Seq data. We identified 3,189 and 3,022 genes that show allele-specific expression (*p* < 0.05) in replicates one and two, respectively (Figure 1, Table S9). Furthermore, we also identified 862 and 911 genes that would have been falsely identified to have allele-specific expression (false positives) if the copy numbers of SNV allele frequencies were not taken into consideration as well as 446 and 407 genes that would not have been identified (false negatives) in replicates one and two, respectively (Table S10).

### Allele-Specific DNA Methylation

Using the phasing information for HepG2 SNVs (Dataset 2), we also identified 384 CpG islands (CGIs) that exhibit allele-specific DNA methylation (Figure 1, Table S11). We obtained two independent replicates of HepG2 whole-genome bisulfite sequencing data (2×125 bp, experiment ENCSR881XOU) from the ENCODE Portal (Sloan et al., 2016). Read alignment to hg19 was performed using Bismark (Krueger and Andrews, 2011); 70.0% of reads were uniquely aligned and a striking 44.7% of cytosines were methylated in a CpG context. We then phased methylated and unmethylated CpGs to their respective haplotypes by identifying reads that overlap both CpGs and phased heterozygous SNVs (Dataset 2). We grouped the phased individual CpGs into CGIs and totaled the number of reads that contain methylated and unmethylated cytosines for each CGI allele, normalizing by CN in cases of aneuploidy. Fisher’s exact test was used to evaluate allele-specific methylation, and significant results were selected using a target false discovery rate of 10% (Storey and Tibshirani, 2003) (see Methods). In total, 98 CGIs reside within promoter regions (defined as 1 kb upstream of a gene); 277 are intragenic, and 96 lie within 1 kb downstream of 348 different genes. The following 11 genes are within 1 kb of a differentially methylated CGI and also overlap with the Sanger Cancer Gene Census: *FOXA1*, *GNAS*, *HOXD13*, *PDE4DIP*, *PRDM16*, *PRRX1*, *SALL4*, *STIL*, *TAL1*, and *ZNF331*.

### Allele-Specific CRISPR Targets

We identified 38,551 targets in the HepG2 genome suitable for allele-specific CRSIPR targeting (Figure 1, Table S12). Phased variant sequences (including reverse complement) that differ by >1 bp between the alleles were extracted to identify all possible CRISPR targets by pattern searching for [G, C, or A]N_20_GG (see Methods). Only conserved high-quality targets were retained by using a selection method previously described and validated (Sunagawa et al., 2016). We took the high-quality target filtering process further by taking the gRNA function and structure into account. Targets with multiple exact matches, extreme GC fractions, and those with TTTT sequence (which might disrupt the secondary structure of gRNA) were removed. Furthermore, we used the Vienna RNA-fold package (Lorenz et al., 2011) to identify gRNA secondary structure and eliminated targets for which the stem loop structure for Cas9 recognition is not able to form (Nishimasu et al., 2014). Finally, we calculated the off-target risk score using the tool as described for this purpose (Ran et al., 2013). A very strict off-target threshold score was chosen in which candidates with a score below 75 were rejected to ensure that all targets are as reliable and as specific as possible.

### Genomic Sequence and Structural Context Provides Insight into Regulatory Complexity

We show examples of how deeper insights into gene regulation and regulatory complexity can be obtained by integrating genomic sequence and structural contexts with functional genomics and epigenomics data (Figure 5A-D). One example is the allele-specific RNA expression and allele-specific DNA methylation in HepG2 at the *PLK2* locus on chromosome 5 (Figure 5A). By incorporating the genomic context in which *PLK2* is expressed in HepG2 cells, we see that *PLK2* RNA is only expressed from Haplotype 1 (*p* = 4.66E-10) in which the CGI within the gene is completely unmethylated (*p* = 1.51E-66) in the expressed allele and completely methylated in the non-expressed allele (Figure 5A, C, D). The second example is allele-specific RNA expression and allele-specific DNA methylation of the *TBX2* gene in HepG2 (Figure 5B). The *TBX2* locus on chromosome 17 is triploid, and we see that *TBX2* is preferentially expressed from Haplotype 1 which has one copy and lower expression is observed from the two copies of Haplotype 2 (*p* = 0.0179) (Figure 5B, C). We also observed highly preferential DNA methylation of the CGI in Haplotype 1 (*p* = 1.55E-32) (Figure 5B, D). In addition, there is also an allele-specific CRISPR targeting site for both haplotypes in the promoter region of *TBX2* and inside CGI 22251 (1,937 bp upstream of *TBX2* gene and 2,259 bp downstream of the 5’ end of CGI 22251) (Figure 5B).

## DISCUSSION

When the HepG2 cell line was first established in 1979, it was mistakenly reported as of hepatocellular carcinoma origin (Aden et al., 1979) and also curated such in the ATCC repository (Rockville, MD, USA). This misclassification has generated much confusion among investigators in the past decades and in the published literature. Review of the original tumor specimen by the original investigators firmly established HepG2 to be of epithelial hepatoblastoma origin rather than hepatocellular carcinoma (López-Terrada et al., 2009a). As one of the most widely used cell lines in biomedical research and one of the main cell lines of ENCODE, HepG2’s genomic sequence and structural features have never been characterized in a comprehensive manner beyond its karyotype (Chen et al., 1993; Simon et al., 1982) and SNVs identified from ChIP-Seq data and 10x coverage WGS that do not take aneuploidy or CN into consideration (Cavalli et al., 2016; Huang and Ovcharenko, 2015). Studies conducted using the extensive collection of functional genomics and epigenomics datasets for HepG2 have previously relied on the human reference genome. Here, we present the first global and phased characterization of the HepG2 genome. By using deep short-insert WGS, 3 kb-insert mate-pair sequencing, array analysis, karyotyping, deep linked-reads sequencing, and integrating a collection of novel and established computational methods, we performed a comprehensive analysis of genomic structural features (Figure 1) for the HepG2 cell line that includes SNVs (Dataset 1), Indels (Dataset 1), large CN or ploidy changes across chromosomal regions at 10 kb resolution (Table S2), phased haplotype blocks (Dataset 2), phased CRISPR targets (Table S12), novel retrotransposon insertions (Table S8), and SVs (Dataset 3, Dataset 4, Dataset 5, Dataset 6) including deletions, duplications, inversions, translocations, and those that are the result of complex genomic rearrangements. Many of the HepG2 SVs are also phased, assembled, and experimentally verified (Dataset 5, Table S7, and Table S8).

We also illustrate, using *PLK2* and *TBX2* (Figure 5A, B), examples where knowing the genomic sequence and structural context can enhance the interpretation of function genomics and epigenomics data to derive novel insights into the complexity of oncogene regulation. The Polo-like kinase gene *PLK2* (*SNK*) is a transcriptional target of p53 and also a cancer biomarker (Burns et al., 2003; Coley et al., 2012). It has been studied in the contexts of many human cancers (Burns et al., 2003; Ou et al., 2016; Pellegrino et al., 2010; Syed et al., 2011). Disruption of PLK2 has also been proposed to have therapeutic value in sensitizing chemo-resistant tumors. Its roles in Burkitt’s lymphoma (Syed et al., 2006), hepatocellular carcinoma (Pellegrino et al., 2010), and epithelial ovarian cancer (Syed et al., 2011) are consistent with that of tumor suppressors while its role in colorectal cancer is consistent with that of an oncogene (Ou et al., 2016). Interestingly, promoter methylation and/or LOH were linked to the down-regulation of PLK2 in human hepatocellular carcinoma (Pellegrino et al., 2010). Chemotherapy resistance of epithelial ovarian cancer can be conferred by the down-regulation of PLK2 at the transcriptional level via DNA methylation of the CpG island in the *PLK2* promoter (Syed et al., 2011). Here we show that the down-regulation of PLK2 in HepG2 cancer cells could be achieved through what appears to be allele-specific transcriptional silencing via allele-specific DNA methylation of a large CGI within the gene body (Figure 5A).

The T-box transcription factor TBX2 is a critical regulator of cell fate decisions, cell migration, and morphogenesis in the development of many organs (Cho et al., 2011; Harrelson et al., 2004; Manning et al., 2006; Suzuki et al., 2005). It regulates cell cycle progression (Bilican and Goding, 2006), and its overexpression has been demonstrated in promoting or maintaining the proliferation of many cancers including melanomas (Vance et al., 2005), nasopharyngeal cancer (Lv et al., 2017), breast cancer (D’Costa et al., 2014; Wang et al., 2012), prostate cancer (Du et al., 2017), and gastric cancer (Yu et al., 2015). Here, we show that three copies of the *TBX2* gene exist in HepG2 cancer cells as a result of duplication in Haplotype 2. However, it is preferentially expressed in Haplotype 1 possibly due to the highly allele-specific DNA methylation in the CGI that span its promoter region and most of the gene body (Figure 5B). It is plausible that overexpression of TBX2 in other cancer types are caused by similar genomic rearrangements and/or epigenetic mechanisms where duplication of *TBX2* may result in the overexpression and DNA methylation (possibly allele-specific) may contribute an additive effect to TBX2 overexpression or act as the sole contributor where *TBX2* is not duplicated.

The duplication of *TBX2* is a direct consequence of aneuploidy. Previous studies have also revealed the aneuploid karyotype of the HepG2 cell line (Chen et al., 1993; Simon et al., 1982). The karyotype of HepG2 cells in our analysis was largely supported by previously published karyotypes for all chromosomes (Table S1). While karyotypes provide a general guide of the degree of aneuploidy in HepG2 cells, we point researchers to the results of our high-resolution read depth analysis of large CN changes across the HepG2 genome for a clearer picture (Table S2). See ***Supplementary Discussion*** for detailed discussion of other oncogenes, tumor-suppressors, and other genes associated with cancer that are disrupted as a consequence of genomic variation in HepG2.

The identification of SNVs and Indels with sensitivity and accuracy requires deep WGS coverage (>33x for SNVs and >60x for Indels) (Bentley et al., 2008; Fang et al., 2014). From our WGS dataset (>70.3x coverage) we identified large numbers of SNVs and Indels that we subsequently corrected for their allele frequencies according to chromosomal CN. In addition to being essential for haplotype discovery, correct allele frequencies of variants are also needed for functional genomics or epigenomics studies such as the identification of allele-specific gene expression or of allele-specific transcription factor binding in HepG2. A statistically significant increase in transcription factor binding signal for ChIP-Seq or transcription signal in RNA-Seq for one allele compared to the other at a heterozygous locus may be seen as a case of allele-specific expression or allele-specific transcription factor binding which usually indicates allele-specific gene regulation at this locus. However, if aneuploidy can be taken into account and the RNA-Seq or ChIP-Seq signals are normalized by CN, the case observed might simply due to increased CN at that particular locus rather than the preferential activation of one allele over the other. This was often the case in our re-analysis of two replicates of HepG2 RNA-Seq data where we identified 862 and 911 genes that would have been falsely identified to have allele-specific expression in addition to 446 and 407 genes that would not have been identified to have allele-specific expression in replicates one and two, respectively, if chromosomal CN or haplotype allele frequency was not taken into consideration (Table S10).

The haplotype phase of the genomic variants is an essential aspect of human genetics but entirely ignored by current standard WGS approaches. To obtain haplotype phasing information for HepG2 (Dataset 2), we performed deep coverage linked-read sequencing (Zheng et al., 2016). However, due to high LOH (Figure 1 and Table S3), chromosomes 22 and large portions of 6, 11, and 14 were difficult to phase resulting in much shorter haplotype blocks compared to other chromosomes (Dataset 2, Figure 1, Figure S4). We further extended on the phasing capabilities of Long Ranger and constructed mega-haplotypes that encompass entire HepG2 chromosome arms (Figure 2E, Table 1, Supplementary Data). This was achieved by leveraging the inherent haplotype imbalance in aneuploid genomic regions in cancer genomes using a recently published method developed specifically for this purpose (Bell et al., 2017). Heterozygous loci in aneuploidy regions that contain more than two haplotypes were excluded from phasing due to software limitations (Zheng et al., 2016). The phase information of these loci could be resolved, in principle, from our linked-read data should new algorithms become available.

Combining orthogonal methods and signals greatly improves SV-calling sensitivity and accuracy (Layer et al., 2014; Mohiyuddin et al., 2015). Here, we combined deep short-insert WGS, mate-pair sequencing, and linked-read sequencing with a combination of several computational SV calling methods to identify a spectrum of structural variants that includes deletions, duplications, and inversions as well as CN-corrected complex rearrangements. We compared SVs identified from various methods. For deletions, we see significant overlap as well as variant calls that are specific to each method (Figure S5A), but overlap is less for duplications (Figure S5B) and inversions (Figure S5C). This is consistent with what has been shown previously (Lam et al., 2012) as inversions and duplications are more difficult in principle to accurately resolve. Since each SV detection method is designed to use different types of signals and also optimized to identify different classes of SVs, such overlaps are also expected. Experimental validation of individual SVs of interest should be conducted prior to functional follow up studies.

All data and results generated from this global whole-genome analysis of HepG2 is publicly available on the ENCODE portal (Sloan et al., 2016). This analysis serves as a valuable reference for further understanding the vast amount of existing HepG2 ENCODE data, such as determining whether a known or potential sequence regulatory element has been altered by SNVs, Indels, Alu or LINE1 insertions, CN changes of that given element, or subjected to allele-specific regulation.

Our results also guide future study designs that utilize HepG2 cells including CRISPR experiments where knowledge of the phased genomic variants can extend or modify the number of editing targets including those that are allele-specific (Table S12) while knowledge of aberrant chromosomal CN changes will allow for more accurate interpretation of functional data in non-diploid regions. This study on the HepG2 genome may serve as a technical archetype for advanced, integrated, and global analysis of genomic sequence and structural variants for other widely cell lines with complex genomes.

Researchers should consider that HepG2 and other widely utilized cell lines have been passaged for long periods of time across many different laboratories and encounter opportunities for additional genome variation to occur, especially if they are HepG2 cells that are long separated from the ENCODE HepG2 production line we used in our analysis. Many of the analyses we discuss here are supported by previous study, for instance, our karyotyping and reported mutation in *CTNNB1*, but there are minor differences such as the lack of a mutation in *CAPRIN2* which has been previously reported. We expect that the vast majority of genomic variants that we describe here can be found across the different versions of HepG2 cells but it is always possible that distinct lines of HepG2 cells may harbor slight variations. This also applies when analyzing the various functional genomic datasets available for HepG2 on the ENCODE Portal (Sloan et al., 2016) that have accumulated over the years, especially if follow up work for individual loci is conducted on a different HepG2 line. A first step should always be to experimentally confirm the presence of the particular genomic variant of interest in that particular working line of HepG2. For global analyses that integrate multi-omics datasets, we expect the majority of the genomic variants described and catalogued here to exist: datasets presented here should be well-suited for global perspectives and substantial insights can be expected to be gained. While the aneuploidy in HepG2 renders the design and interpretation of HepG2 genomic and epigenomic studies more challenging, the results of this study enables researchers to continue to use HepG2 to investigate the effects of different types of genomic variations on the multiple layers of functionality and regulation for which ENCODE data is already available and continues to be produced. Although our studies reveal that the genome of HepG2 highly complex, taking our results into account for future analyses of HepG2 ENCODE data should not be considered to make the process more challenging but rather potentially much more insightful and rewarding. This study is primarily focused on the utilization of Illumina sequencing to resolve the genome structure of HepG2. In the future, we plan to utilize other long-read technologies such as Pacific Biosciences and Oxford Nanopore.

## METHODS

To characterize the aneuploid genome of HepG2 in a comprehensive manner, we integrated karyotyping (Figure S1), high-density oligonucleotide arrays, deep (70.3x non-duplicate coverage) short-insert whole-genome sequencing (WGS), 3kb-mate-pair sequencing, and 10X Genomics linked-reads sequencing (Zheng et al., 2016). Using read-depth analysis of WGS data (Abyzov et al., 2011), we first obtained a high-resolution aneuploid map or large copy number (CN) changes by chromosomal region. CN changes were also validated by karyotyping and two independent replicates of arrays. This high-resolution aneuploid map was then integrated into the identification of SNVs and indels and in determining their heterozygous allele-frequencies. By leveraging the inherent haplotype imbalance in aneuploid genomic regions, SNV haplotype blocks were “stitched” into mega-haplotypes that encompass entire chromosome arms (Bell et al., 2017). In addition, SVs, such as deletions, duplications, inversions, and non-reference retrotransposon (LINE1 and Alu) insertions, were also identified from the deep-coverage WGS dataset. The 10X Genomics linked-reads dataset was used to phase haplotypes as well as to identify, assemble, and phase primarily large (>30kb) SVs that include translocations (Marks et al., 2018; Spies et al., 2017; Zheng et al., 2016). The 3kb-mate pair was used to validate the large SVs resolved from using the linked-read sequencing data and also to identify additional SVs, mostly in the medium size-range (1kb-100kb). Finally, we used the phased haplotype information to produce an allele-specific CRISPR targeting map for HepG2 and identified cases of allele-specific expression and allele-specific DNA methylation. See Supplementary Methods for detailed descriptions.

## DATA AVAILABILITY

Raw and processed data files are publicly released on the ENCODE portal (encodeproject.org) via experimental accession numbers ENCSR350RUO, ENCSR276ECO, and ENCSR319QHO. Datasets 1-6 can be accessed via ENCODE accession numbers ENCFF336CFC, ENCFF853HHD, ENCFF467ETN, ENCFF717TPE, ENCFF330UFT, ENCFF241CEK respectively. For immediate review of Datasets and Supplementary Data, files are also available under https://stanfordmedicine.box.com/s/w1fa3ncuhje8zabgugvsw6wmeu7ojv3k

## AUTHORS’ CONTRIBUTIONS

B.Z. and A.E.U conceived and designed the study. B.Z., R.P., I.S., and Y.H. performed experiments. B.Z., S.S.H., S.U.G., N.S., J.M.B., XL.Z., XW.Z., J.G.A., S.B., G.S., D.P., and A.A. performed analysis. H.P.J, W.H.W., and A.E.U contributed resources and supervised the study. B.Z., S.S.H., and A.E.U. wrote the manuscript.

## ACKNOWLEDGEMENTS

We thank Arineh Khechaduri for performing genomic DNA preparation. Aditi Narayanan, Dr. Idan Gabdank, Nathaniel Watson, Zachary A. Myers, and Dr. Cricket Sloan for data organization and upload to ENCODE. Dr. Athena Cherry and the Stanford Cytogenetics Laboratory for karyotype analysis. A.E.U. was supported by NIH grant HG007735 and the Stanford Medicine Faculty Innovation Program. W.H.W. received support from NIH grants HG007834 and HG007735. B.Z. was additionally funded by NIH Grant T32 HL110952. J.G.A was funded by the NSF Graduate Research Fellowship and NIH T32-GM096982.

---

**Figure S1.**
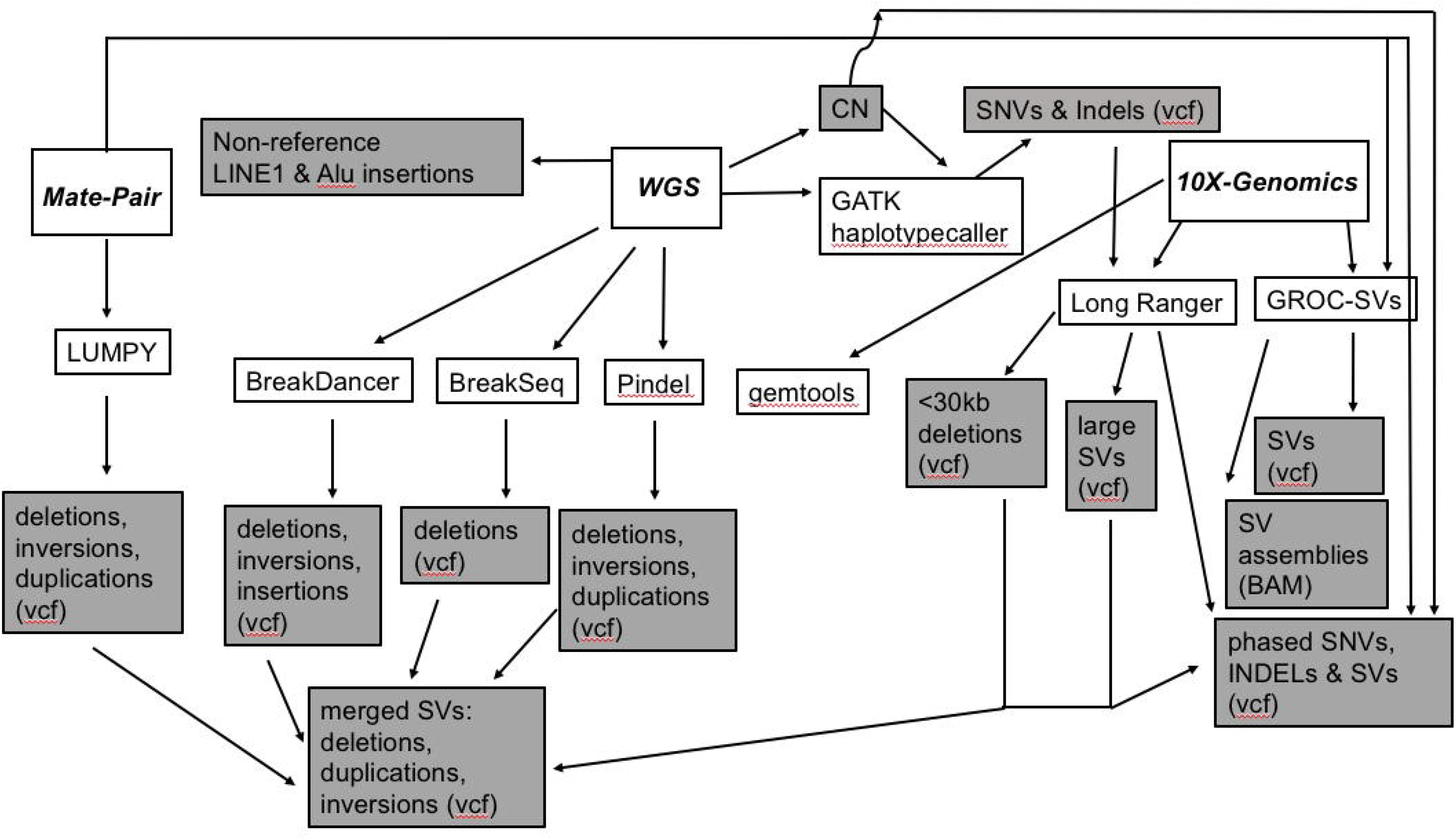
Illustration of the Sequencing-Based Methodologies Used. Deep short-insert WGS (70x non-duplicate coverage), 3 kb-mate-pair sequencing (Korbel et al., 2007), and 10X-Genomics linked-reads sequencing (Zheng et al., 2016) were used to comprehensively characterize the genome of HepG2. The WGS dataset was used to obtain CN i.e. ploidy by chromosome segments using read-depth analysis (Abyzov et al., 2011), SNVs and Indels using GATK Haplotypecaller with CN taken into account (McKenna et al., 2010), non-reference LINE-1 and Alu insertions (Keane et al., 2013), and SVs, such as deletions, duplications, inversions, and insertions using BreakDancer (Chen et al., 2009), BreakSeq (Lam et al., 2010), and Pindel (Ye et al., 2009). The linked-read sequencing data was used to phase heterozygous SNVs and Indels as well as to identify, assemble, and phase large (>30 kb) and SVs using Long Ranger (Zheng et al., 2016) and GROC-SVs (Spies et al., 2017). Deletions <30 kb were also identified by Long Ranger. The SV assembly file from GROC-SVs is in BAM format (Dataset 5). The 3 kb-mate-pair sequencing data was used to call additional structural variants, mostly in the medium size-range (1 kb-100 kb) using LUMPY (Layer et al., 2014) and also used to validate large and complex SVs. The union of non-complex SVs identified merged into a single VCF file (Supplementary Data).

**Figure S2.**
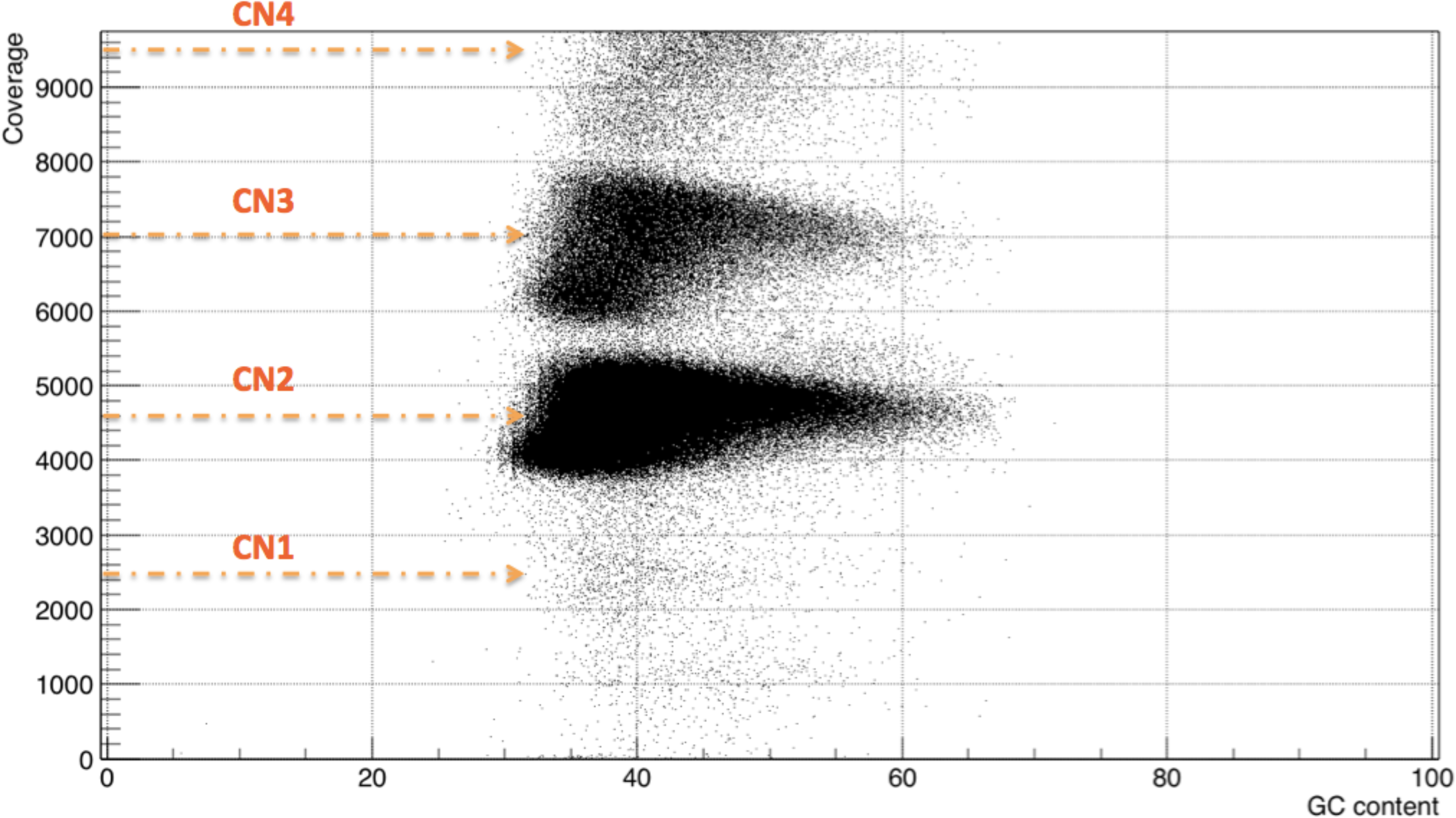
HepG2 WGS coverage vs. % GC content. Y-axis: HepG2 short-insert WGS coverage in 10 kb bins across the genome; X-axis: % GC content of bins. coverage). Clusters correspond to CN (i.e. ploidy).

**Figure S3.**
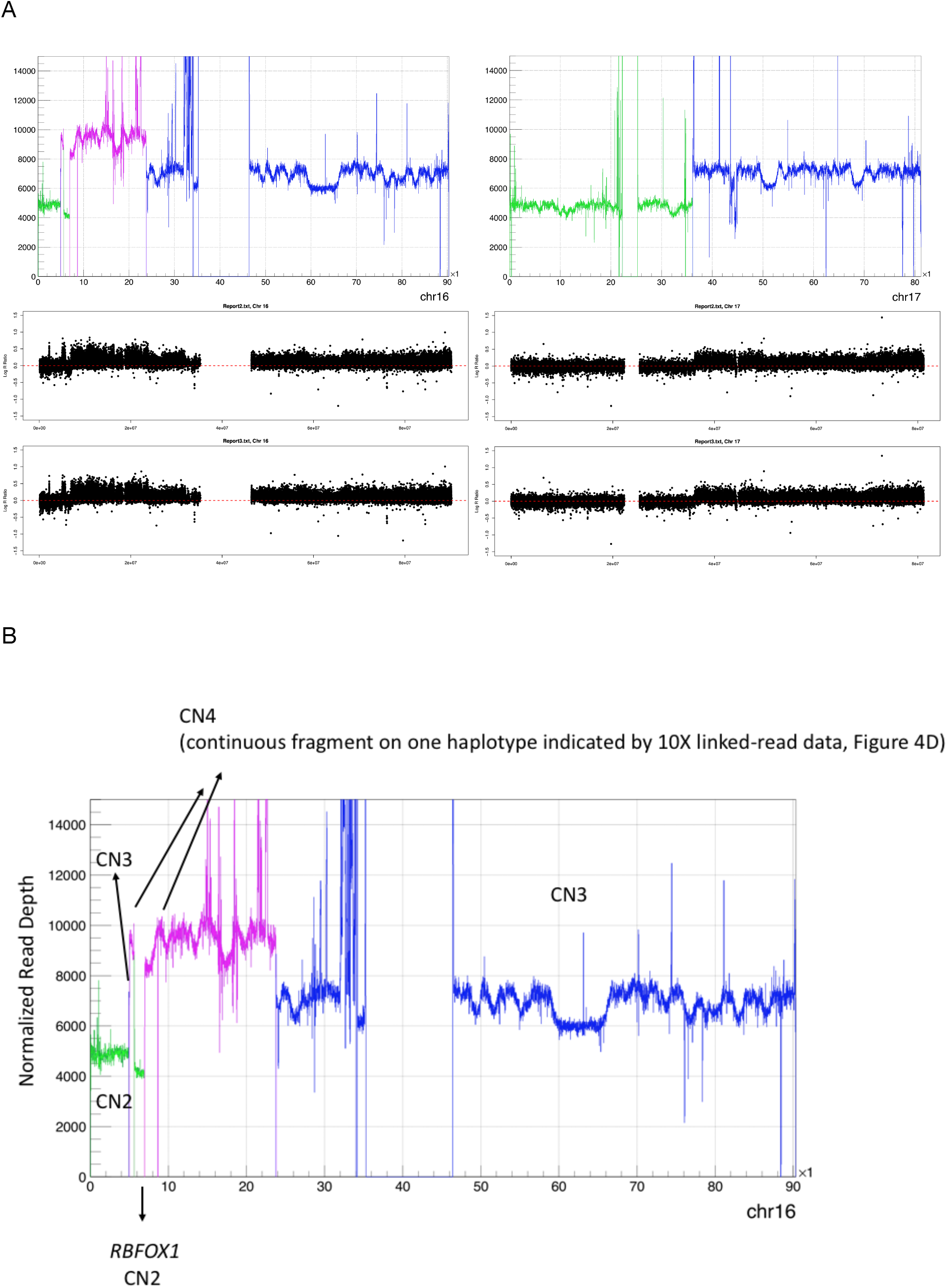
Orthogonal Validation of CN as Determined by Read-Depth Analysis in HepG2 by Illumina MEGA Array (2 Replicates) (Chromosomes 16 & 17) (A) Upper panel: WGS coverage plot. X-axis genomic coordinate in kb. Y-axis: WGS coverage. Purple: CN4, Blue: CN3, Green: CN2. Lower panel: Y-axis: array probe signal intensity. X-axis: Genomic coordinate. For complete chromosomes (1-22, X), see Supplementary Data. (B) Copy numbers of genomic regions on chromosome 16 of HepG2. X-axis genomic coordinate in kb. Y-axis: WGS coverage. Purple: CN4, Blue: CN3, Green: CN2.

**Figure S4.**
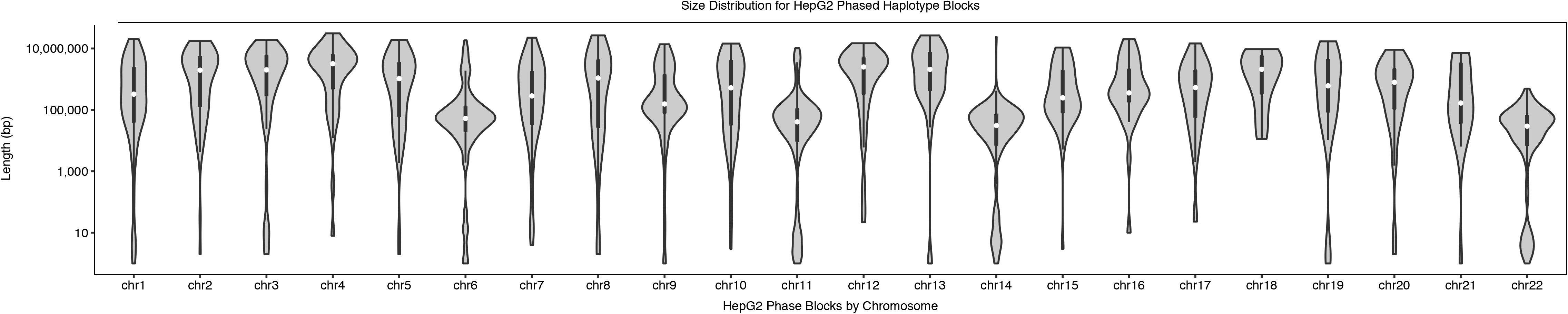
Size Distribution of Phased Haplotype Blocks by Chromosome. Violin plots (with overlaid boxplot) of HepG2 phased haplotype blocks (Dataset 2) by chromosomes. Y-axis: size in log-scale.

**Figure S5.**
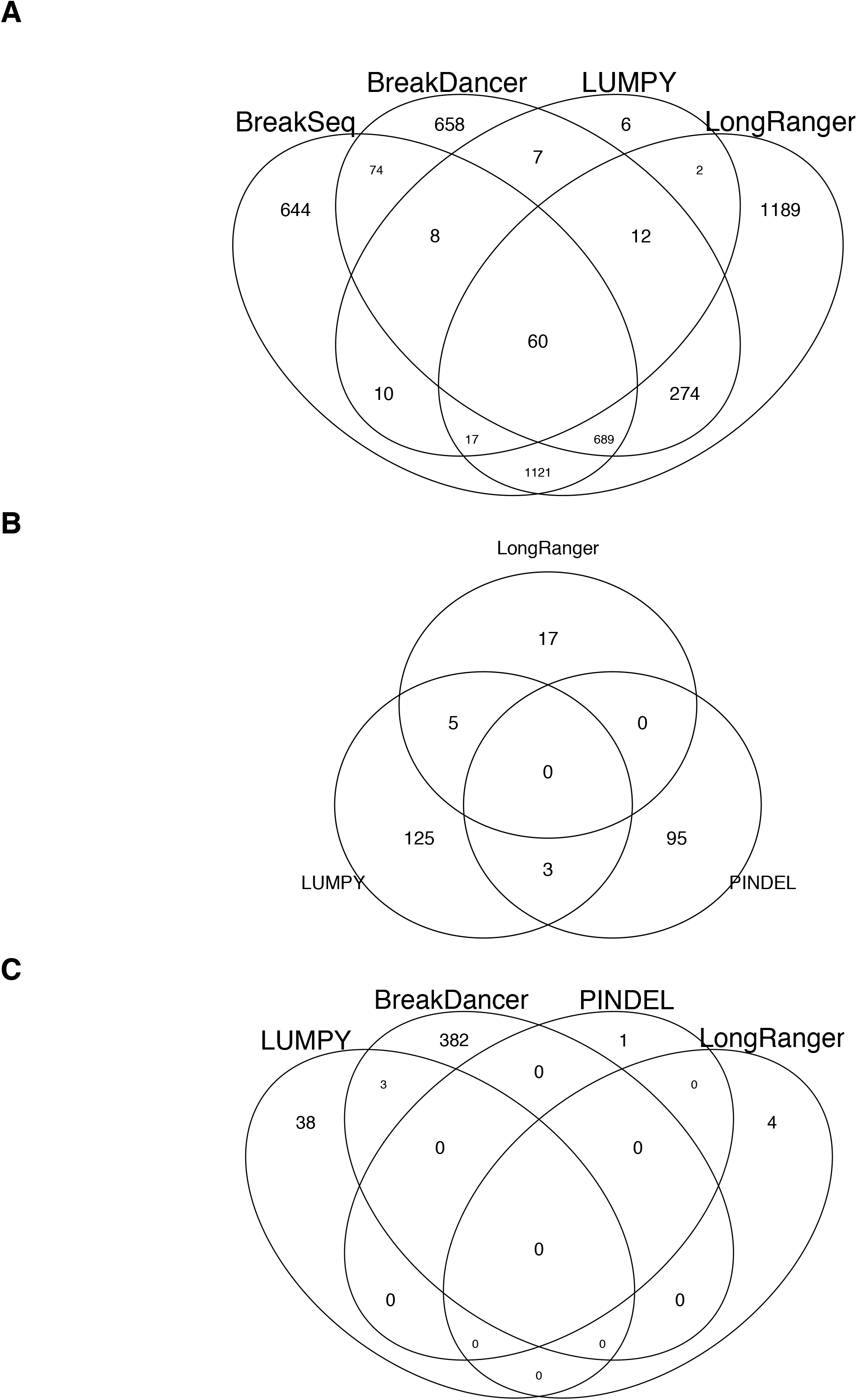
Overlap Between SV Callers. Venn diagram of overlaps (>50% reciprocal) between HepG2 SVs identified in WGS using BreakDancer (Chen et al., 2009), Pindel (Ye et al., 2009), and BreakSeq (Lam et al., 2010), in 3kb-mate-pair sequencing using LUMPY (Layer et al., 2014), and in linked-read sequencing using Long Ranger (Marks et al., 2018; Zheng et al., 2016) for (A) deletions (>50 bp), (B) tandem duplications, and (C) inversions.

